# A Novel Model of Estrogen Receptor-Positive Breast Cancer Bone Metastasis with Antiestrogen Responsiveness

**DOI:** 10.1101/2021.09.16.460665

**Authors:** Kendall L. Langsten, Lihong Shi, Adam S. Wilson, Brian Westwood, Maria T. Xie, Victoria E. Surratt, JoLyn Turner, Ravi Singh, Katherine L. Cook, Bethany A. Kerr

## Abstract

Most women diagnosed with breast cancer (BC) have estrogen receptor alpha positive (ER+) disease. ER+ BC preferentially metastasizes to bone; at which time it is considered incurable. Treatments for bone metastasis have not advanced in decades, in part due to a lack of appropriate ER+ BC bone metastasis models. We developed an immunocompetent ER+ BC murine model with spontaneous bone metastasis and antiestrogen responsiveness. To do this, we transduced triple-negative (TN) bone-tropic murine BC cell lines 4T1.2 and E0771/Bone to express ERα. These cells were then injected into the mammary fat pads of Balb/c (n=21) or C57Bl/6 (n=27), respectively. Once tumors established, mice were treated with either the selective estrogen receptor modulator (SERM) tamoxifen (TAM), the selective estrogen receptor degrader (SERD) ICI 182,780 (ICI, Faslodex, fulvestrant), or vehicle control for 21 days. Tumor volumes and weights significantly decreased in the ER+ groups treated with TAM and ICI compared with ER+ vehicle-treated groups. Staining for immune profiles and total RNA sequencing demonstrated modified immune cell infiltration between TN and ER-derived tumors. Approximately 25% of the mice with ER+ 4T1.2 tumors developed metastases to long bones while none of the mice with TN tumors developed metastases. This immunocompetent ER+ 4T1.2 BC model may allow for further exploration of ER+ BC bone metastasis mechanisms and for the development of new therapeutics for women diagnosed with bone metastasis from ER+ BC.

**Simple Summary:** Estrogen receptor alpha positive (ER+) breast cancer is the most common subtype of breast cancer. When it metastasizes to bone, it becomes incurable. Little advancement has occurred in the treatment of bone metastasis from ER+ breast cancer, partly due to the lack of animal models. To establish an animal model of ER+ BC, we genetically modified two triple-negative breast cancer cell lines to express ERα and injected the cell lines into murine mammary glands. Mice were treated with standard antiestrogen therapies, the selective estrogen receptor modulator tamoxifen or the selective estrogen receptor degrader ICI 182,780. We found that compared to mice with triple-negative breast cancer, mice with ER+ breast cancer developed bone metastases and were responsive to antiestrogen therapy. This model allows for further exploration of bone metastasis mechanisms and for the development of new therapeutics, translating into improved clinical outcomes for women with bone metastasis from ER+ breast cancer.

## 1. Introduction

Breast cancer (BC) is the most common non-skin related cancer diagnosed in women globally and is the leading cause of female cancer mortality in 110 countries [1]. Cancer associated deaths are overwhelmingly related to metastasis [2] and of the women with metastatic BC, 65-80% will have bone metastases [3]. While early diagnosis of BC prior to bone metastasis is ideal, approximately 3.7% of women with BC will have bone metastases at the time of first diagnosis [4]. Once in the bone, metastases are associated with extreme pain, bone fractures, nerve, and spinal compression, and are considered incurable [5]. Better strategies are needed to prevent and treat bone metastases of BC.

A significant factor in the development of bone metastasis is positive estrogen receptor (ER+) status in the breast tumor. ER+ BC is the most common subtype of BC representing approximately 70% of all BC cases [6,7]. While ER+ BC bone metastases presents a substantial clinical problem, few advancements have been made for the prevention and treatment of disease in recent decades [8]. This is due, in part, to the fact that there are few animal models of ER+ BC that metastasize to the bone spontaneously after injection into the mammary gland. Currently, most models of ER+ BC metastasis rely on intra-cardiac injection of tumor cells [9–11]. While this does result in tumor growth in many tissues, including bone, it leaves researchers without the ability to study the initial steps of the metastatic cascade where tumor cells in the mammary gland infiltrate surrounding tissue and gain access to the vasculature. Studies of BC metastasis also often rely on exogenous estrogen which modifies the bone microenvironment, meaning that it is not entirely representative of human disease [12]. Models of BC that originate in the mammary gland and metastasize to bone currently utilize triple-negative (TN) BC cell lines, and many use genetically modified or immunocompromised mouse strains, which can be difficult to maintain and may not translate well to human disease [13–16]. A recent study demonstrated bone metastasis after intraductal injection of ER+ ZR751 human xenograft cells [17]; however, the xenograft-based system did not allow for analysis of interaction with the immune system. Until researchers have a model of ER+ BC that spontaneously metastasizes to bone from the mammary gland in an immunocompetent environment, mechanisms driving the initial metastatic cascade and the tendency towards metastasizing to bone cannot be fully understood.

Considering the above limitations, we developed a model of ER+ BC that spontaneously metastasizes to the bone from the mammary gland and is responsive to antiestrogen therapy in immunocompetent mice. Furthermore, we characterized the immune profiles, RNA profiles, and metastatic ability of the ER+ tumors. We genetically modified two bone-tropic, TN BC cell lines to express ERα and injected the cells into the mammary glands of mice. Mice were then treated with commonly used antiestrogen therapies, either the selective estrogen receptor modulator (SERM) tamoxifen (TAM) or the selective estrogen receptor degrader (SERD) ICI 182,780 (ICI). Using multiple methods (histology, immunohistochemistry, and RNA sequencing), we found that when compared to mice with TN tumors, ER+ tumors were larger and were responsive to antiestrogen therapy. Furthermore, there were differences in immune cell infiltration within TN and ER+ tumors. Only mice with ER+ 4T1.2 tumors developed bone metastases. This immunocompetent ER+ 4T1.2 model holds the potential to direct advancement in treatment for the underserved population of women with ER+ BC bone metastasis.

## 2. Materials and Methods

### 2.1 Cell Line Generation

The E0771/Bone and 4T1.2 murine bone-tropic, TNBC lines were provided under material transfer agreements with Drs. Hiraga and Anderson, respectively [14,16]. These two sublines were derived from the parental TNBC 4T1 and E0771 to have increased bone tropism by repeated intracardiac injection and isolation of resulting bone metastatic subclones. Using an ERα-GFP construct developed by Dr. Elaine Alarid, we generated ER+ sublines of both cell lines in collaboration with the Wake Forest Comprehensive Cancer Center Cell Engineering Shared Resource (CESR). To confirm its identity, the ERα-GFP construct was sent to GeneWiz (South Plainfield, NJ) for Sanger sequencing using standard M13-F and M13-R sequencing primers. The ERα-GFP expression cassette was cloned into a lentivirus transfer vector using methods based upon the ViraPower™ Lentiviral Gateway™ Expression system (Thermo Fisher Scientific, Waltham, MA). Briefly, following sequencing, the ERα-GFP expression cassette was PCR amplified to introduce the attB and attL-R sites using following primers:

Forward: 5’- CACCACGGCCACGGACCATGA - 3’

Reverse: 5’- TTACTTGTACAGCTCGTCCATGCCGAG - 3’

A Gateway entry clone was generated using the Gateway pENTR/D-TOPO linear vector cloning kit (Thermo Fisher Scientific) to insert the ERα-GFP expression cassette into the Gateway donor vector. Restriction analysis at Noti-HF (NEB #R3189L, 20K U/mL, Lot 10030794), NruI-HF (NEB #R3192S, 20K U/mL, Lot 10030601), and AgeI-HF (NEB #R3552S, 20K U/mL, Lot 10028839) was performed with a CutSmart Buffer at 37°C for either 5-15min or overnight to verify the correct orientation of the insert. The ERα-GFP expression cassette was then transferred from the entry clone to a Gateway compatible lentivirus transfer vector (pLenti CMV Blast DEST (706-1), a gift from Eric Campeau & Paul Kaufman (Addgene plasmid #17451; http://n2t.net/addgene:17451; RRID:Addgene_17451, [18]) using LR Clonase II. The transfer vector was then packaged into a lentivirus using the pPACKH1 HIV third generation lentiviral expression system kit and PureFection reagent (both from System Biosciences, Palo Alto, CA) into HEK293-T cells (ATCC, Manassas, VA). Lentiviral particles were concentrated using the Clontech Lenti-X Concentrator (Takara Bio USA, Mountain View, CA). The concentrated lentivirus was used to transduce the cell lines, and ERα-GFP expressing cells were obtained after antibiotic selection for two weeks. A lentivirus transfer vector expressing GFP alone (pLenti CMV GFP Puro (658-5), a gift from Eric Campeau & Paul Kaufman (Addgene plasmid #17448; http://n2t.net/addgene:17448; RRID:Addgene_17448, [18]) was packaged into lentiviral particles as described above. Both TNBC parental cell lines were transfected with this vector and GFP expressing cells obtained after antibiotic selection. All cells tested negative for mycoplasma and murine viruses prior to injection into animals.

### 2.2 Animal experiments

All experiments were performed in accordance with the Institutional Animal Care and Use Committee at the Wake Forest School of Medicine (IACUC A20-10). Seven-week-old, intact female, Balb/c (n=21) and C57Bl/6 (n=27) mice from Jackson Laboratory (Bar Harbor, MI) were housed with 12-hour dark/light cycles and ad libitum access to food and water. Mice were restrained and 1×105 TN or ER+ 4T1.2 or E0771/Bone cells suspended in 20 μL of sterile phosphate buffered saline (PBS) were injected into the 4th inguinal mammary fat pad. Tumors were allowed to establish and reach a volume of 100 mm3. At this time, mice with ERα expressing tumors were treated with either tamoxifen (TAM) at 5 mg over 30 days as a time release pellet (ER+ 4T1.2, n=6; ER+ E0771/Bone, n=7), ICI 182,780 (ICI) at 1 mg per week subcutaneously (ER+ 4T1.2, n=6; ER+ E0771/Bone, n=7), or a subcutaneous injection of sterile PBS as a vehicle control (ER+ 4T1.2, n=4; ER+ E0771/Bone, n=6). Tumor volumes were measured every three days. After 21 days of treatment, mice were humanely euthanized and tumors, visceral organs, and long bones were collected. Tumors were weighed and collected fresh-frozen or in 4% paraformaldehyde prior to being embedded in paraffin. After fixation in 4% paraformaldehyde, the hind limb bones were isolated from whole legs and decalcified in 14% neutral buffered EDTA for 2 weeks prior to being embedded in paraffin.

### 2.3 Immunohistochemistry

Immunohistochemistry on formalin-fixed paraffin-embedded tumors and hind limbs from mouse experiments were performed to characterize the tumors and identify metastatic tumor cells. Paraffin embedded tumors and bones were sectioned at 5 μm thick and placed on charged glass slides. For tumors, antigen unmasking was performed by heat-induced epitope retrieval using 0.05% citraconic anhydride solution (pH 7.4) for 45 minutes at 98°C; for bone samples, antigen unmasking was performed by heat-induced epitope retrieval using Tris-EDTA PH 9.0 for 20 minutes at 95°C. Endogenous horseradish peroxidase (HRP) activity was quenched by incubating with BLOXALL blocking solution for 10 minutes. Samples were blocked with 1% BSA for 30 minutes at room temperature then incubated with primary antibody overnight at 4°C. Immunohistochemistry of the tissue sections was performed using antibodies against CD3, CD68, CD45R, and neutrophil elastase on the primary tumors and GFP, pan-cytokeratin, Sca-1, and endomucin on the bone (Table 1). After incubation, samples were washed with PBS then incubated with HRP-conjugated secondary antibody (Vector Laboratories, Burlingame, CA) followed by Nova Red chromogen (Vector Laboratories, Burlingame, CA) for staining development. All samples were counterstained with hematoxylin. Slides were then scanned using a Hamamatsu NanoZoomer by the Virtual Microscopy Core in the Wake Forest School of Medicine. Immunostaining was quantified using the VisioPharm digital pathology analysis software. Briefly, the total area of region of interest (ROI) was measured for each specimen; custom-designed apps were then used to identify and measure regions with positive staining within the ROI, and the ratios of the positive staining area vs the total area were calculated. Three separate, non-consecutive tissue sections of whole tibiae (including trabecular regions in the metaphysis, the epiphysis, and the diaphysis) were analyzed for pan-cytokeratin or GFP staining to quantify numbers of metastases in the tibiae.

**Table 1.** Primary antibodies used for immunohistochemistry of animal primary tumors and bone.

| Antibody | Concentration | Company | Catalogue No. |
| --- | --- | --- | --- |
| CD3 | 1:750 | Abcam | Ab16669 |
| CD68 | 1:2000 | Abcam | Ab125212 |
| CD45R (B220) | 1:500 | Abcam | Ab64100 |
| Neutrophil elastase | 1:500 | Abcam | Ab68672 |
| GFP | 1:1000 | Thermo Fisher Scientific | OSE00003G |
| Pan-cytokeratin | 1:1000 | Cell Signaling Technology | 4545S |
| Sca-1 | 1:400 | Abcam | Ab109211 |
| Endomucin | 1:200 | Thermo Fisher Scientific | 14-5851-82 |

### 2.4 Bone Structure and Osteoclast Analysis

To visualize changes in the bone structure and osteoclasts, paraffin-embedded bones were sectioned at 5 μm and placed on charged glass slides. Sections were stained for 1 hour at 37oC in tartrate-resistant acid phosphatase (TRAP) staining solution (0.3 mg/mL Fast Red Violet LB, 0.05 M sodium acetate, 0.03 M sodium tartrate, 0.05 M acetic acid, 0.1 mg/mL naphthol, 0.1% Triton X-100, pH 5.0) to visualize osteoclasts. Sections were counterstained with Gill Method Hematoxylin Stain 1 (ThermoFisher). Slides were scanned at 20x with the Hamamatsu NanoZoomer by the Virtual Microscopy Core in the Wake Forest School of Medicine. Bone histomorphometry, osteoclast numbers, and growth plate organization were analyzed with the BioQuant Osteo software (RRID: SCR_016423) using a ROI approximately 150 μm distal to the metaphysis encompassing the trabecular region of the tibiae. Osteoclasts are detected based on thresholding for the red/purple TRAP staining.

### 2.5 RNA sequencing and Gene Set Enrichment

To extract RNA for sequencing, sections of TN and ER+ 4T1.2 derived tumors were collected on ice and washed in PBS at a pH of 7.4 until all blood and debris was removed. Total RNA was extracted using the QIAGEN RNeasy mini-Kit (Qiagen GMBH, Hilden, Germany) per the manufacturer’s protocol. The concentration of total RNA was estimated using a Nanodrop One C (Thermo Fisher Science) and RNA library formation was performed using the Illumina HiSeq 6000 (Illumina), S4 flowcell platform with PE150 seq parameter by Novogene. To mine for gene sets that were enriched in the ER+ tumors, datasets provided by Novogene (Ensembl output) were filtered using both membership in SwissProt and Entrez conversion in DAVID [19–21]. RNA log-fold change was only considered in the analysis if the p-value was <0.1 and there was an average count of 64 or more for all subjects in the analysis. GSEA pre-ranked analysis [22,23] was run for KEGG pathways, collapsed to gene symbol for filtered set. If a pathway was identified with an FDR≤ 0.25, then genes in that pathway were considered enriched. Enriched gene sets in the mouse tumors were analyzed in ER+ and ER- tumors from BC patients using the publicly available METABRIC (Molecular Taxonomy of Breast Cancer International Consortium) dataset [24] via cBioPortal (www.cbioportal.org). Briefly, tumors with ER positivity and negativity were queried and compared, each of the genes identified in the mouse gene set were searched and the fold change was calculated using cBioPortal. Genes that were significantly dysregulated from the human gene set identified via cBioPortal were plotted against patient survival for ER+ BC patients using KMplotter (http://kmplot.com) [25,26].

### 2.6 Statistical Analysis

To determine statistical significance, Student’s t test, one-way, or two-way analysis of variance (ANOVA) with Tukey post-test were used to analyze data with the GraphPad Prism 9 software (RRID: SCR_002798). Error bars represent the SEM of experiments. * p<0.05, ** p<0.01, and *** p<0.005.

## 3. Results

### 3.1 ER+ tumors were larger than TN tumors and were responsive to antiestrogen treatments

To examine the effects of ER-reexpression and antiestrogen treatment on tumor growth, bone-tropic, BC 4T1.2 or E0771/Bone cells with and without ER were injected into Balb/c or C57Bl/6 mammary fat pads, respectively. These two sublines demonstrate increased bone tropism after derivation by repeated intracardiac injection and isolation of bone metastatic clones from the TNBC 4T1 or E0771 parental cell lines [14,16]. Tumor volumes were significantly elevated (approximately 1.5-fold increase) for ER+ 4T1.2 (900 mm3 average) and ER+ E0771/Bone (1,677 mm3 average) derived tumors when compared with the tumors derived from TN parental cell lines (658 mm3 and 1,029 mm3 average, respectively; Two-way ANOVA, p<0.05, Figure 1A, 1C). At study termination, average tumor weight for ER+ E0771/Bone (2.44 g average) was significantly heavier (1.7-fold increase) than TN E0771/Bone (1.17 g average; Figure 1B, One-way ANOVA, p<0.05). Additionally, ER+ 4T1.2 tumors weighed more (1.4-fold increase; 0.98 g average) when compared with TN 4T1.2 tumors (0.69 g average; Figure 1D, One-way ANOVA, p<0.05). All mouse groups with ER+ tumors that were treated with either TAM or ICI had significantly lower tumor volumes and weights than ER+ tumors without treatment (Two-way ANOVA, p<0.05, Figure 1A-D), except for ER+ E0771/Bone treated with TAM which tended to have lower tumor volumes and weights, but was not significantly decreased. These data demonstrate that ER+ expression results in increased tumor growth.

**Figure 1.**
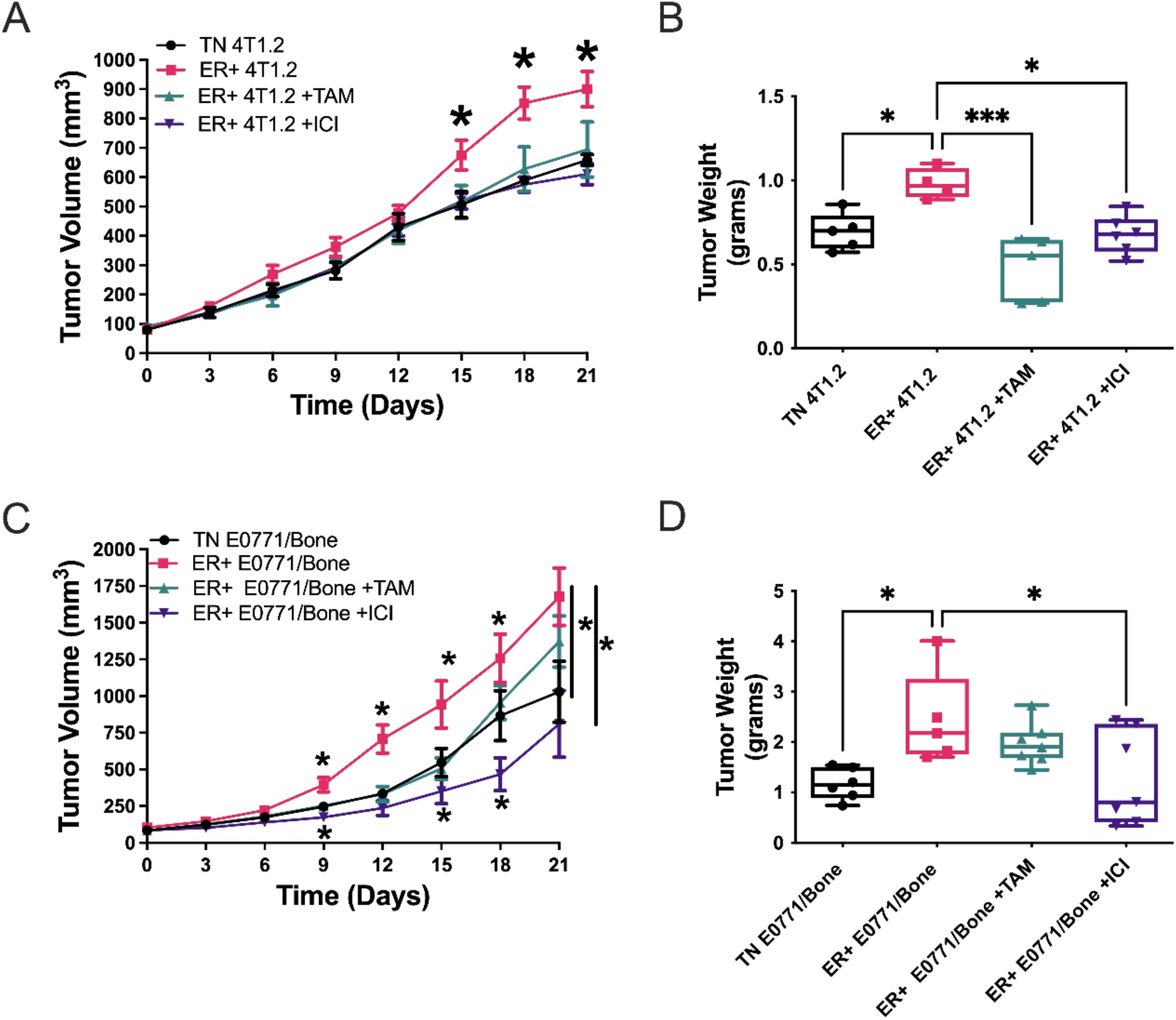
ER+ tumors are significantly larger and responsive to antiestrogen treatment. Parent TN and ER+ 4T1.2 (A and B) or E0771/Bone (C and D) BC cells were injected into the 4th inguinal mammary fat pad in Balb/c or C57Bl/6 mice, respectively. Once tumor volumes reached 100 mm3, mice were treated with vehicle control, 5 mg/ 30-day time-release pellet tamoxifen (TAM), or 1 mg/wk ICI 182,780 (ICI). Tumor volume was tracked over time (A and C) and are represented as mean tumor volume ±SEM (n=4-7). Tumor weight was measured upon experimental termination (B and D) and are represented as mean tumor weight ±SEM (n=4-7). * represents p<0.05 and *** represents p<0.005 by two-way (A and C) or one-way ANOVA (B and D).

### 3.2 ER+ tumors had a modified inflammatory microenvironment compared with TN

The immune landscape within a breast tumor has implications of the patient’s prognosis, most notably with CD68+ macrophage infiltration being associated with worse prognosis [27]. By examining ER+ tumor growth in immunocompetent animals, we can better define the immune landscape in ER+ and TN BC. Immunohistochemistry for immune cells were performed on TN and ER+ tumors to measure differences in immune cell infiltration between the tumor types (Figure 2A). In ER+ tumors, while not significantly altered, the percentage of CD3+ T cells were decreased by 0.9-fold and 0.7-fold and CD45R+ B cells were increased by 2.3-fold and 2.0-fold for ER+ 4T1.2 and ER+ E0771/Bone, respectively when compared with TN derived tumors (Figure 2B, 2C, Student’s T-test, p>0.05). The ER+ E0771/Bone tumors had significantly increased percentages of neutrophil elastase positive cells (24.5-fold increase) and CD68+ macrophages (3.5-fold increase) compared with the TN E0771/Bone tumors (Figure 2C, Student’s T-test, p<0.05). The ER+ 4T1.2 tumors displayed a similar but non-significant trend of increased neutrophil and macrophage infiltration (Figure 2B). Thus, neutrophil and macrophage recruitment may be increased in ER+ tumors compared with their TN counterparts.

**Figure 2.**
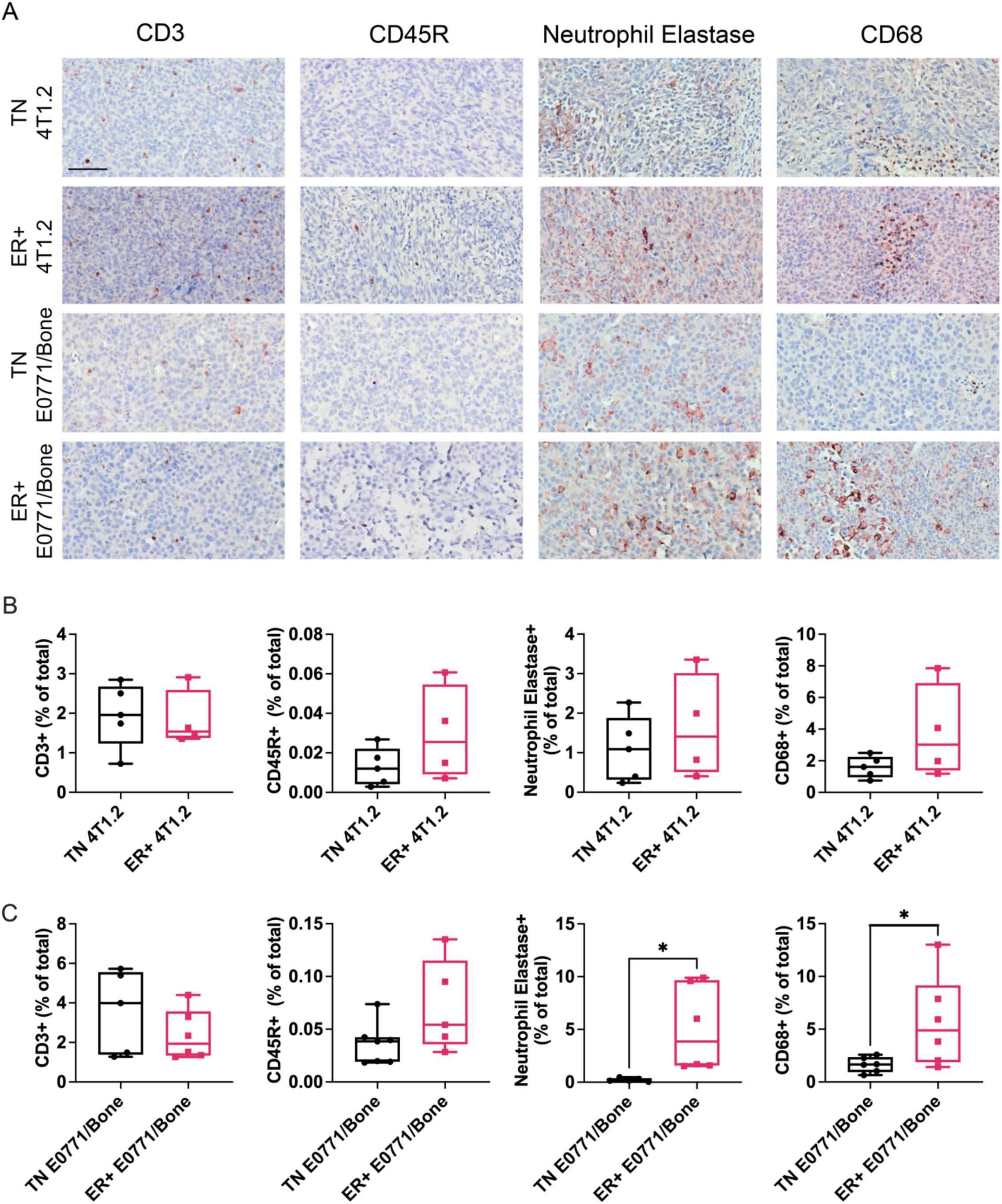
ER+ primary tumors have a modulated immune landscape compared with TN tumors. Representative images of immunohistochemistry (IHC) on primary TN and ER+ 4T1.2 and E0771/Bone (A). Scale bar represents 70 μm. Quantification of tumors for CD3+ T cells, CD45R+ B cells, CD68+ macrophages, and neutrophil elastase (neutrophils). Using the pathology analysis software, Visiopharm, the percentage of immunoreactive cells out of the total tumor area was determined with a custom-made app. Percentages of positive cells are represented as single values with the mean ± SEM for the 4T1.2 TN and ER+ (B) and E0771/Bone TN and ER+ (C) tumors (n=4-7). * represents p<0.05 by Student’s T-test.

### 3.3 ER+ 4T1.2 tumors had significantly downregulated T cell signaling receptor pathways when compared with TN tumors

To further characterize the differences between ER+ and TN BC tumors, we performed RNA sequencing on TN and ER+ 4T1.2 derived tumors. RNA expression profiles have been utilized to determine prognostic factors and potential therapeutic targets for women [28–30]. We identified a 21 gene set signature negatively enriched in ER+ tumors (n=4) compared with TN tumors (n=5, Figure 3A) associated with decreased T cell receptor signaling pathways, indicating a possible mechanism for the slightly decreased in CD3+ T cell infiltration demonstrated in Figure 2. Of those 21 genes, 17 were significant downregulated in the ER+ tumors (Table 2, p<0.05). Using the METABRIC human breast cancer data set [24], the gene set negatively enriched in ER+ mouse tumors was found to largely be significantly downregulated in human ER+ tumors compared with ER negative tumors. Three genes were significantly downregulated in humans only, FYN, CDK4, and PIK3R5 and three genes significantly downregulated in mice only, RASGRP1, NFATC2, and PI3KR1 (Figure 3B; Table 2). Only one of the genes, AKT1, was not significantly decreased in either mice or human ER+ tumors when compared with ER negative, although in both species it tended to be decreased. Of the 14 genes significantly downregulated in both humans with ER+ BC and our mouse model of ER+ BC, expression levels in four were significantly associated with survival outcome (Figure 3C; n=877, p<0.05). Of the four genes, lower expression of ZAP70 (−0.4- and −1.1-fold change in humans and mice, respectively), GRAP2 (−0.1- and −1.4-fold change in humans and mice, respectively), and CD3G (−0.4- and −1.6-fold change in humans and mice, respectively) were associated with worse survival outcomes while lower expression of CARD11 was associated with increased survival outcomes in women with ER+ BC (Figure 3C; n=877, p<0.05). Genes that were significantly downregulated in humans and mice were plotted against survival for ER+ tumors (non-significant Kaplan-Meier curves shown in Figure 4). Our data demonstrate a potential immune gene signature associated with worse prognosis in ER+ BC.

**Figure 3.**
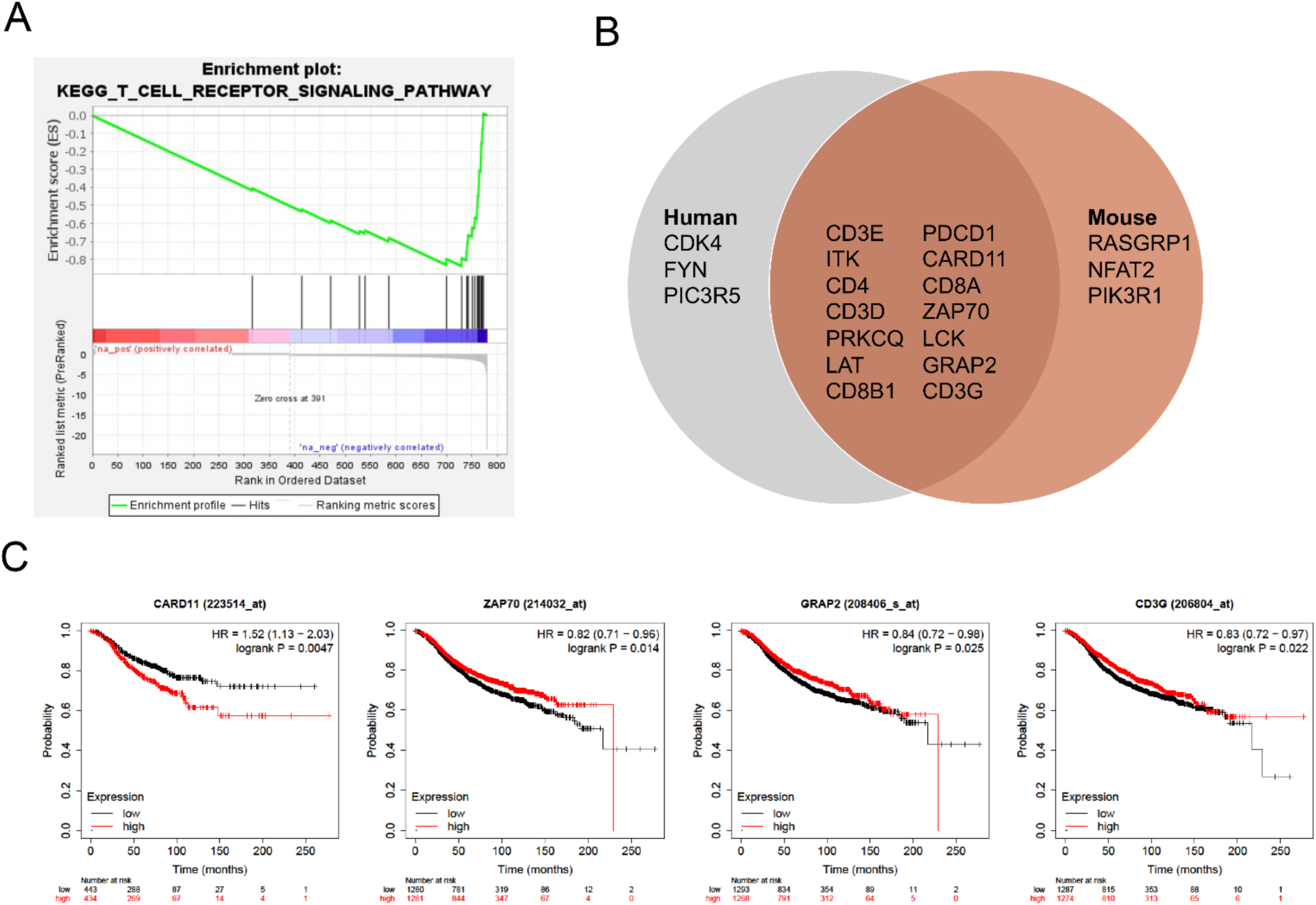
Gene sets associated with T-cell signaling pathways were down regulated in ER+ primary tumors in mice and humans when compared with TN tumors and some of the shared genes were associated with worse outcomes in patients with ER+ tumors. RNA sequencing data from TN 4T1.2 (n=5) and ER+ 4T1.2 (n=4) tumors was performed by Novogene and gene set enrichment was performed using SwissProt keywords in DAVID followed by KEGG pathway analysis. A 21 gene-set involved in T-cell receptor signaling pathways identified with 17 of the genes being significantly negatively enriched in ER+ 4T1.2 tumors compared with TN 4T1.2 tumors (A; p<0.05). To determine clinical significance of the gene set, the negatively enriched gene set was used in cBioPortal to compare human RNA expression data from the METABRIC study. Log fold change for the genes identified in our study were analyzed in ER+ and ER negative tumors, fifteen of the 21 genes were significantly decreased in both ER+ humans and mice with 4T1.2 ER+ tumors (B). Three genes were significantly down regulated in humans only, FYN, CDK4, and PIK3R5 and three genes significantly down regulated in mice only, RASGRP1, NFATC2, and PI3KR1 (B). Of the 14 genes significantly downregulated in both humans with ER+ BC and our mouse model of ER+ BC, expression levels in four were significantly associated with survival outcome (C; n=877, p<0.05). Of the four genes, CARD11, ZAP70, GRAP2, and CD3G, in all but CARD11 lower expression was associated with worse survival.

**Table 2.**
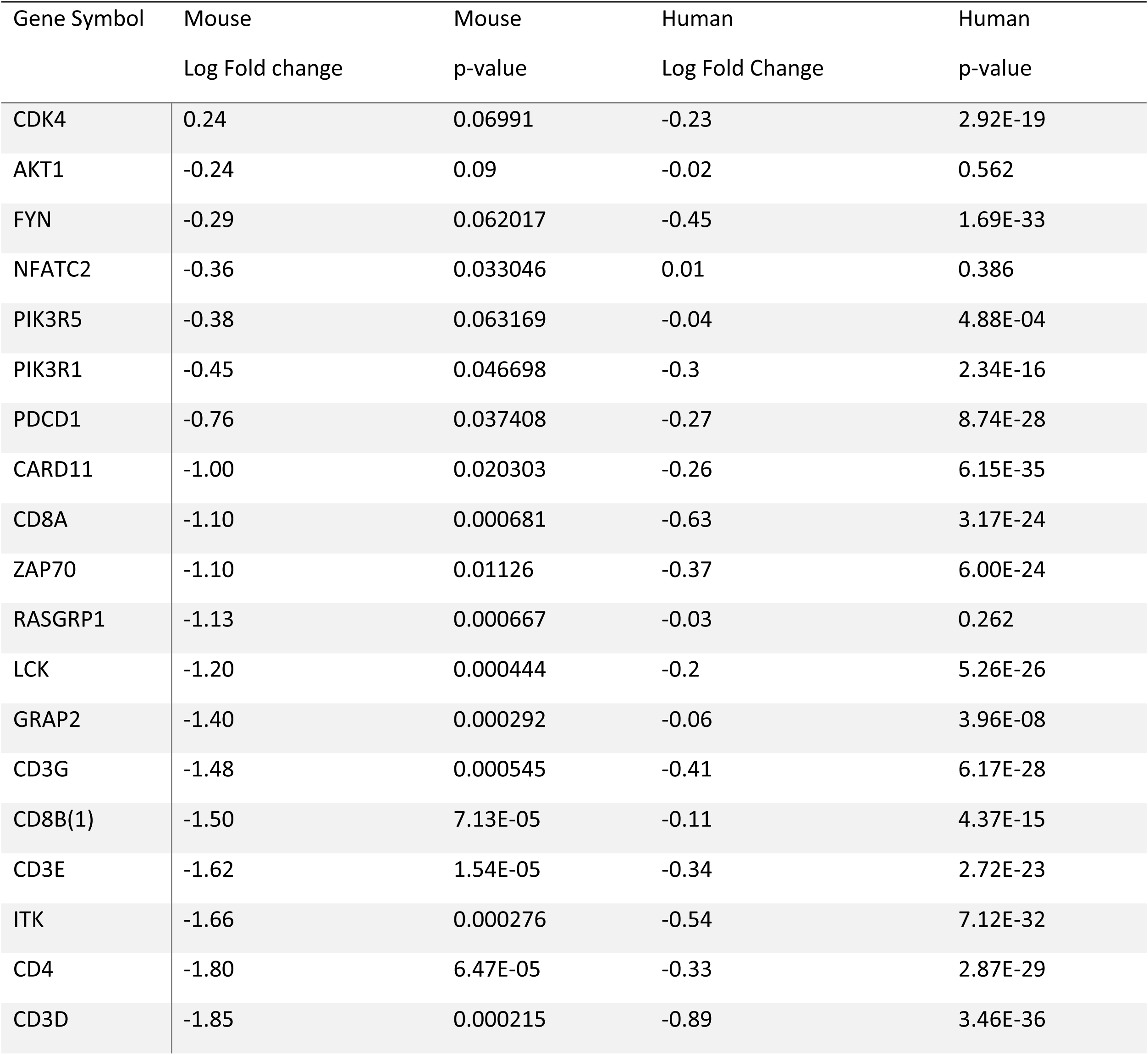
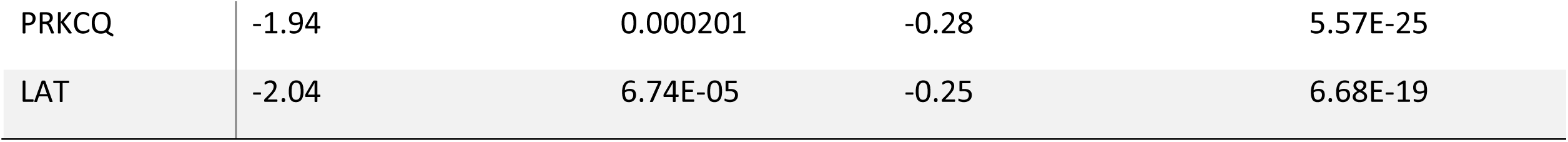
Log fold changes between RNA expression of T cell receptor signaling pathways in TN and ER+ tumors. RNA sequencing on 4T1.2 (n=5) and 4T1.2 ER+ (n=4) tumors from mice was analyzed for enriched gene sets. A set of 21 genes associated with down regulated T cell receptor pathways was downregulated in ER+ tumors compared with TN tumors, with 17 of the 21 being significantly downregulated. RNA expression of these genes was compared between ER+ and ER negative tumors in the METABRIC data set. Gene names, log fold change, and significance level are shown.

**Figure 4.**
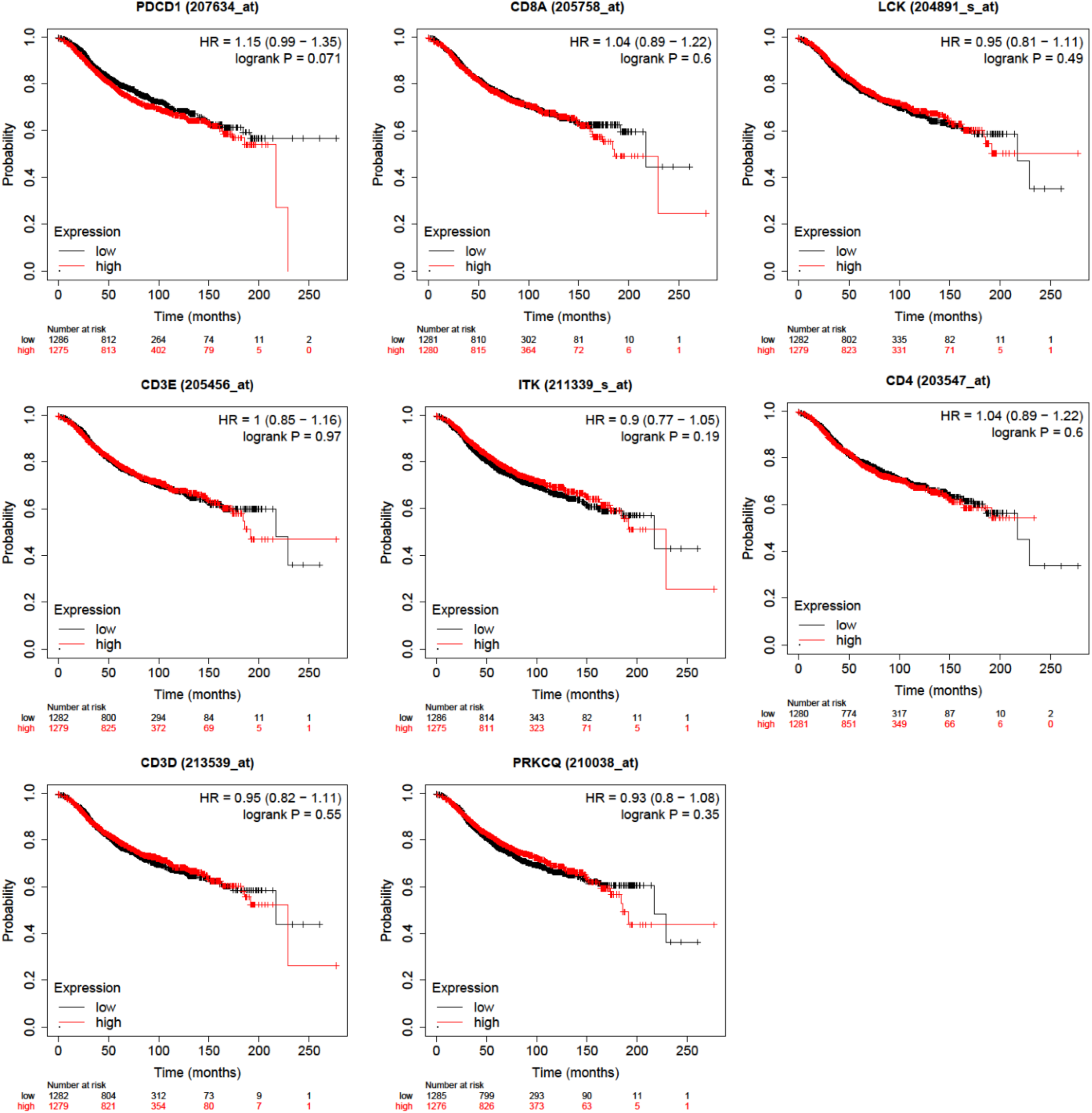
Kaplan-Meier curves of the not significantly downregulated genes in ER+ human tumors from the publicly available METABRIC data source. The expression level of the above genes was not significantly associated with differences in ER+ BC patient survival (n=877, p>0.05).

### 3.4 ERα positive E0771/Bone cells demonstrate potential metastasis to liver

BC demonstrates metastasis to visceral organs such as the lung and liver. The TN 4T1.2 cell line was previously demonstrated to spread to both bone and lungs [16]. Additionally, the parental TN E0771 line demonstrated metastasis to lungs, while the TN E0771/Bone line spread to bone [14]. To test for potential metastatic spread, visceral organ weights were measured at study termination. Lung weights between all treatment groups were not significantly different for TN (0.27 g, 0.16 g) or ER+ (0.26 g, 0.18g) 4T1.2 or E0771/Bone cells, respectively (One-Way ANOVA, p>0.05; Figure 5A,C). Liver weights were 1.2-fold heavier in the ER+ E0771/Bone mice when compared with parental TN line (One-Way ANOVA, p<0.01; Figure 5B). Mice with ER+ E0771/Bone tumors treated with ICI had significantly decreased liver weights when compared with the non-treated ER+ cohort (One-way ANOVA, p<0.05; Figure 5B). Thus, ER expression in E0771/Bone may promote liver metastasis.

**Figure 5.**
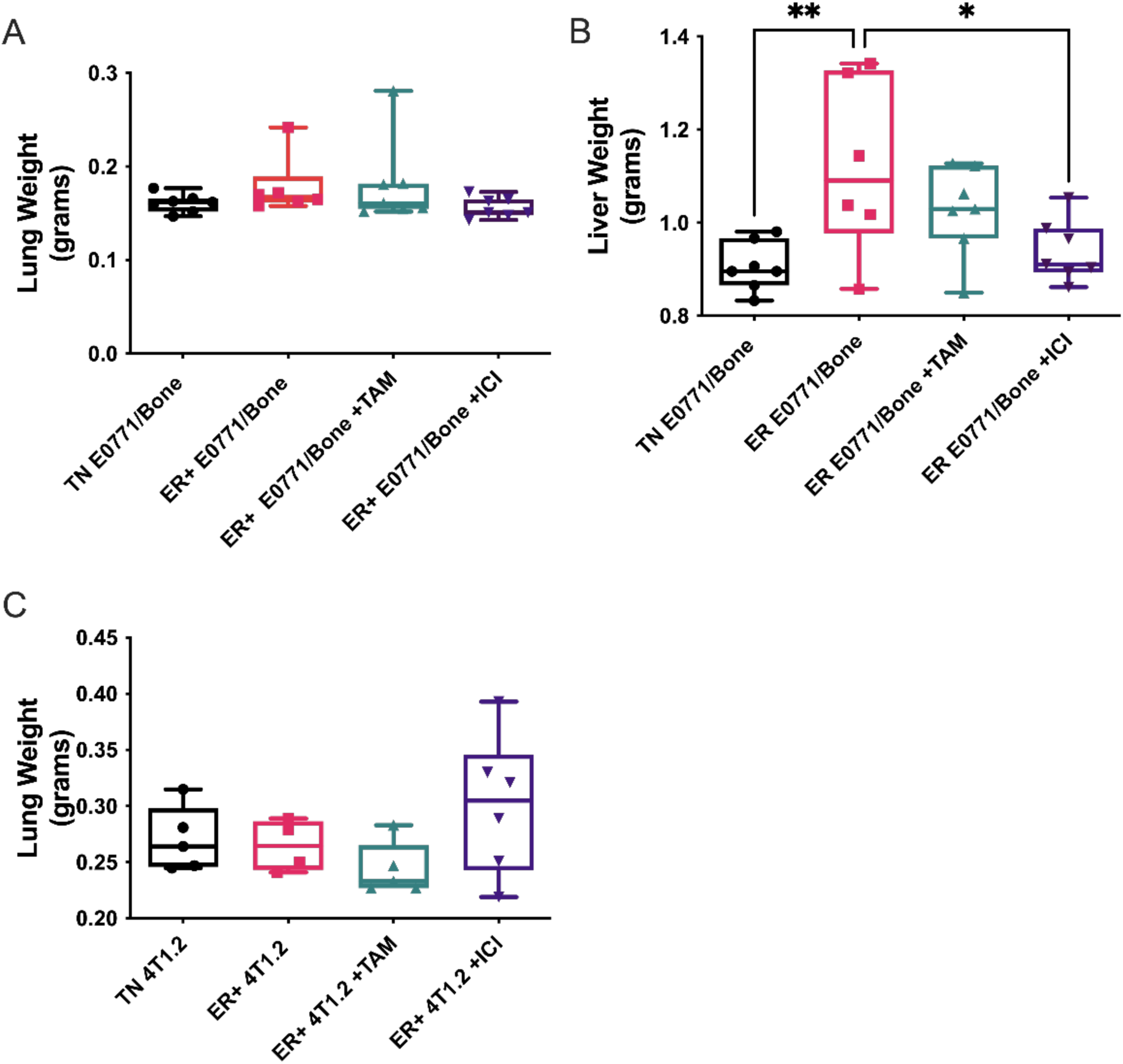
Lung and liver weights representing soft tissue metastases. Lungs and livers were isolated from mice after tumor implantation with E0771/Bone (A and B) or 4T1.2 (C) TNBC or ER+ breast cancer cells and treatment with vehicle controls, 5 mg/ 30 day time release pellet tamoxifen (TAM), or 1 mg/wk ICI 182,780 (ICI). Tissues were weighed and represented as mean weight ± SEM (n=4-7). * represents p<0.05 and ** represents p<0.01 by one-way ANOVA.

### 3.5 Only ERα positive 4T1.2 tumors spontaneously metastasized to bone

To further examine metastatic spread, hindlimbs were isolated from mice injected with ER+ BC after treatment with tamoxifen or ICI and compared to hindlimbs from TN BC injected mice. Tibiae of mice were examined with immunohistochemistry for anti-GFP or anti-pan-cytokeratin, which would identify the GFP labeled TN and ER+ tumor cells injected into the mice and epithelial cells. Only mice injected with ER+ 4T1.2 mice had visible metastases within the bone (Figure 6A). Interestingly, metastases were observed in all ER+ 4T1.2 groups, regardless of treatment (Figure 6B), likely due to the establishment of tumors prior to the initiation of treatment.

**Figure 6.**
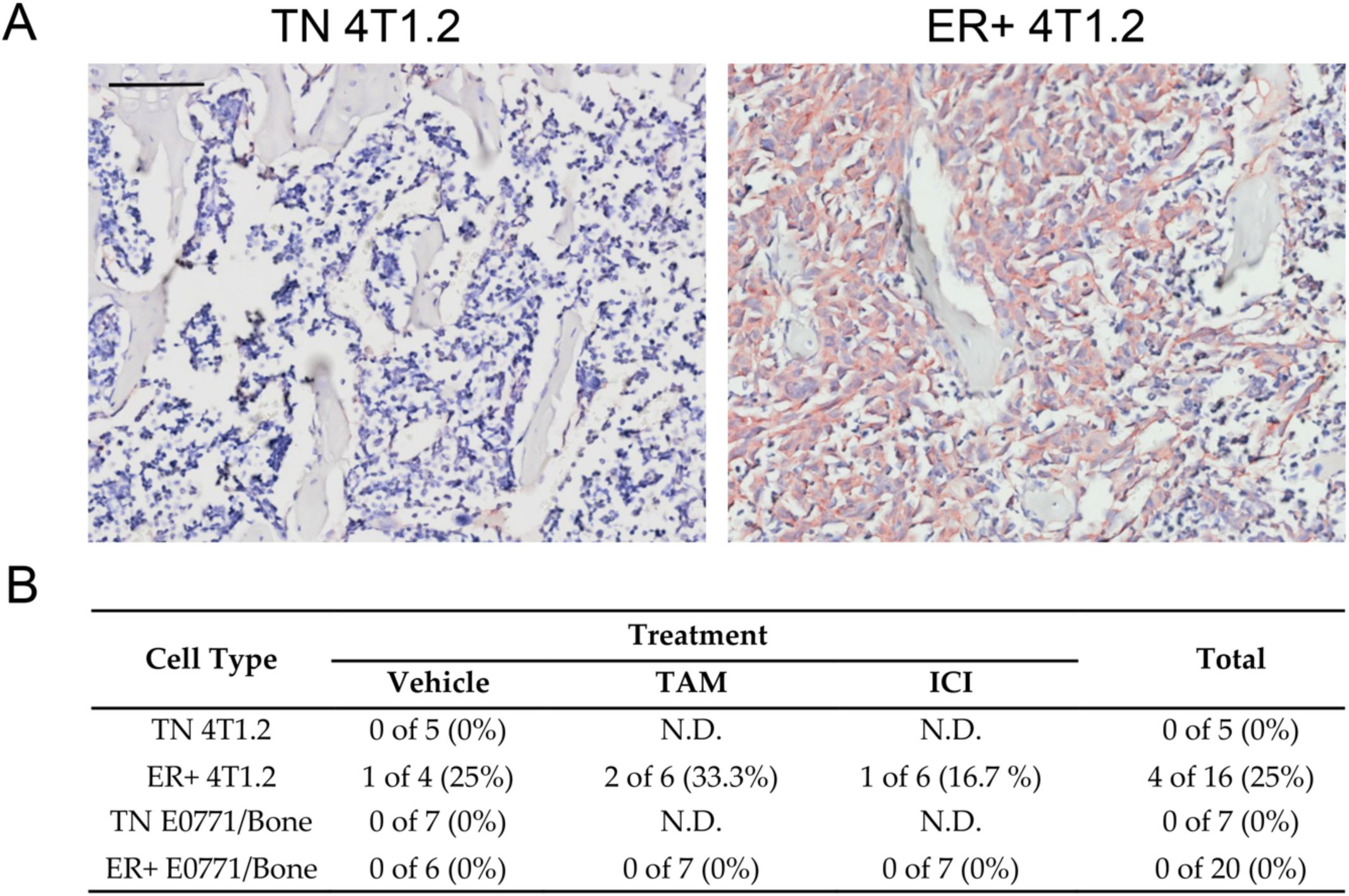
Only ER+ 4T1.2 tumors metastasized to the bone. Immunohistochemistry on tibiae from all mice (n=48) for pan-cytokeratin were performed and analyzed using VisioPharm pathology analysis software. Scale bar represents 150 μm. Only the ER+ 4T1.2 tumors metastasized to the bone, regardless of antiestrogenic treatment. (A) Examples of TN 4T1.2 and ER+ 4T1.2 pan-cytokeratin IHC at the metaphysis are shown. Scale bar represents 150 μm. (B) Numbers of mice with metastatic lesions are quantified with percentages of positive mice shown in the table for each cell type and treatment group. N.D.; not determined

Disseminated BC cells enter the bone microenvironment through the vasculature and colonize the bone at the hematopoietic stem cell (HSC) niche [31]. To assess whether changes in colonization niche were responsible for alterations in bone metastatic spread, we stained bones for Sca-1+ HSC cells and endomucin+ blood vessels (Figure 7A). We found that mice with ER+ E0771/Bone derived tumors had significantly increased percentages of Sca-1 positive HSC (64.4-fold increase) and endomucin positive vasculature (2.8-fold increase) within the bone marrow when compared with TN E0771/Bone tumors (Figure 7C and E). There were no significant differences between Sca-1 or endomucin positivity between mice with TN or ER+ 4T1.2 derived tumors (Figure 7B and D). These data indicate that ER+ 4T1.2 spread to bone was not due to altered colonization niches. These data further suggest that ER+ E0771/Bone tumor growth may induce premetastatic bone microenvironment alterations.

**Figure 7.**
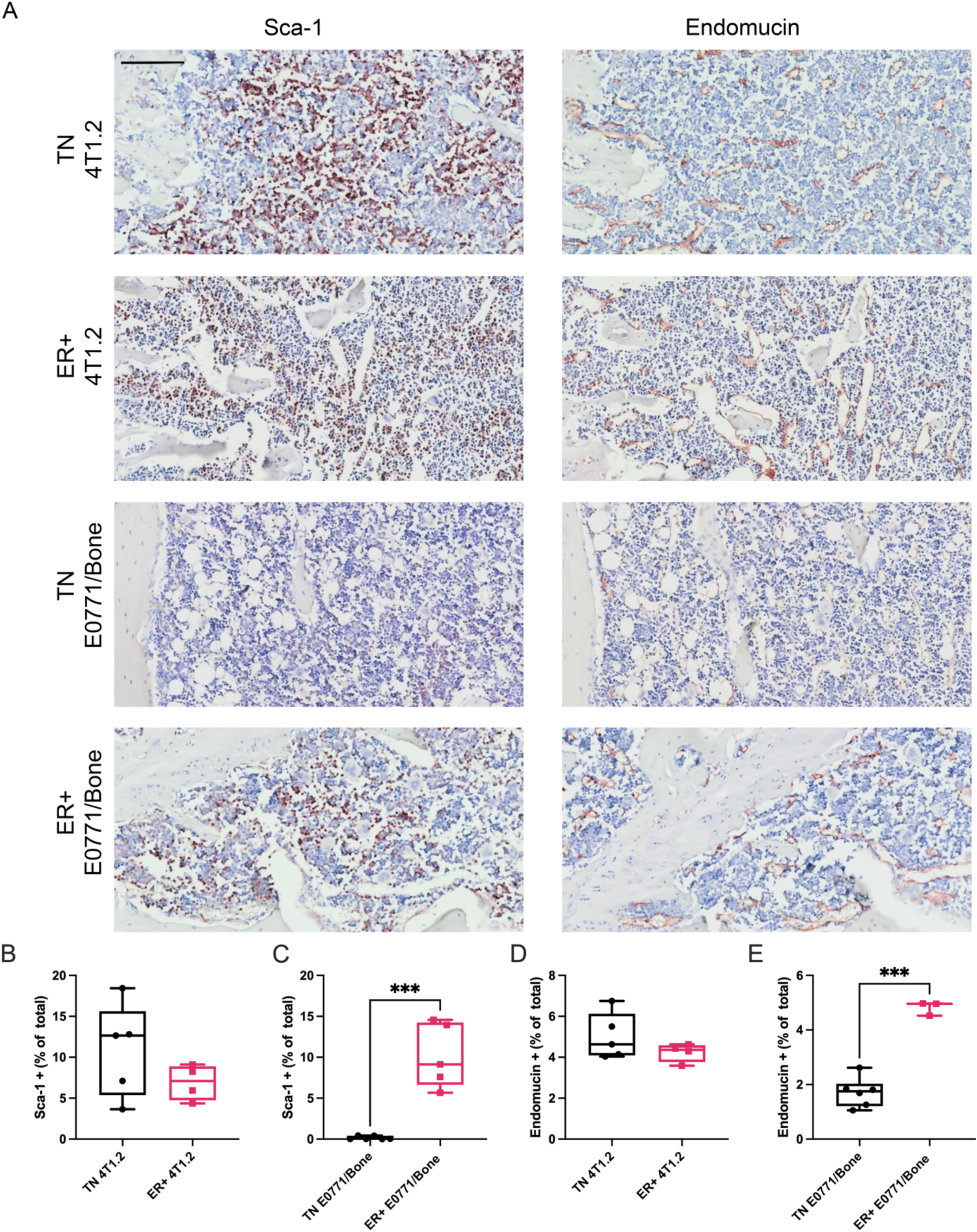
Within the tibiae of mice with ER+ E0771/Bone tumors, there were increased hematopoietic stem cells and vasculature compared with mice with TN E0771/Bone derived tumors. IHC for hemopoietic stem cell marker Sca-1 (A) was performed on tibiae from mice with TN and ER+ 4T1.2 and E0771/Bone tumors and was analyzed using a custom-made app on the pathology analysis software, Visiopharm (n=4-7, B and C). Scale bar represents 150 μm. There was a significant increase in percentage of Sca-1 positive area within the medullary cavity of the mice with ER+ E0771/Bone tumors when compared with mice with TN E0771/Bone (mean ± SEM, n=5-7, Student’s T-test, *** represents p<0.005; C). IHC for the blood vessel marker endomucin (A) was performed on tibiae from mice with TN and ER+ 4T1.2 and E0771/Bone tumors and was analyzed using a custom-made app on the pathology analysis software, Visiopharm (n=4-7, D and E). There was a significant increase in percentage of endomucin positive area within the medullary cavity of the mice with ER+ E0771/Bone tumors when compared with mice with TN E0771/Bone (mean ± SEM, n=3-7, Student’s T-test, *** represents p<0.005; E).

### 3.6 Tamoxifen induced alterations in the premetastatic bone niche

To further characterize premetastatic alterations in the bone microenvironment, we examined the bone structure and osteoclast differentiation. Histomorphometry on tibiae just distal to the growth plate was used to determine the bone volume to tissue volume fraction. No significant changes in the bone fraction was measured between mice with ER+ and TN BC tumors (Figure 8A and C). Mice injected with ER+ 4T1.2 cells that were treated with TAM had significantly increased bone volume: tissue volume percentage when compared with ER+ 4T1.2 cell injected mice that were treated with ICI (Two-way ANOVA, p<0.01, Figure 8A). Similarly, mice injected with ER+ E0771/Bone cells that were treated with TAM tended to have increased bone volume: tissue volume percentage when compared with all other groups (Figure 8C). Thus, TAM treatment alone may alter the bone structure in ways that could alter metastatic spread.

**Figure 8.**
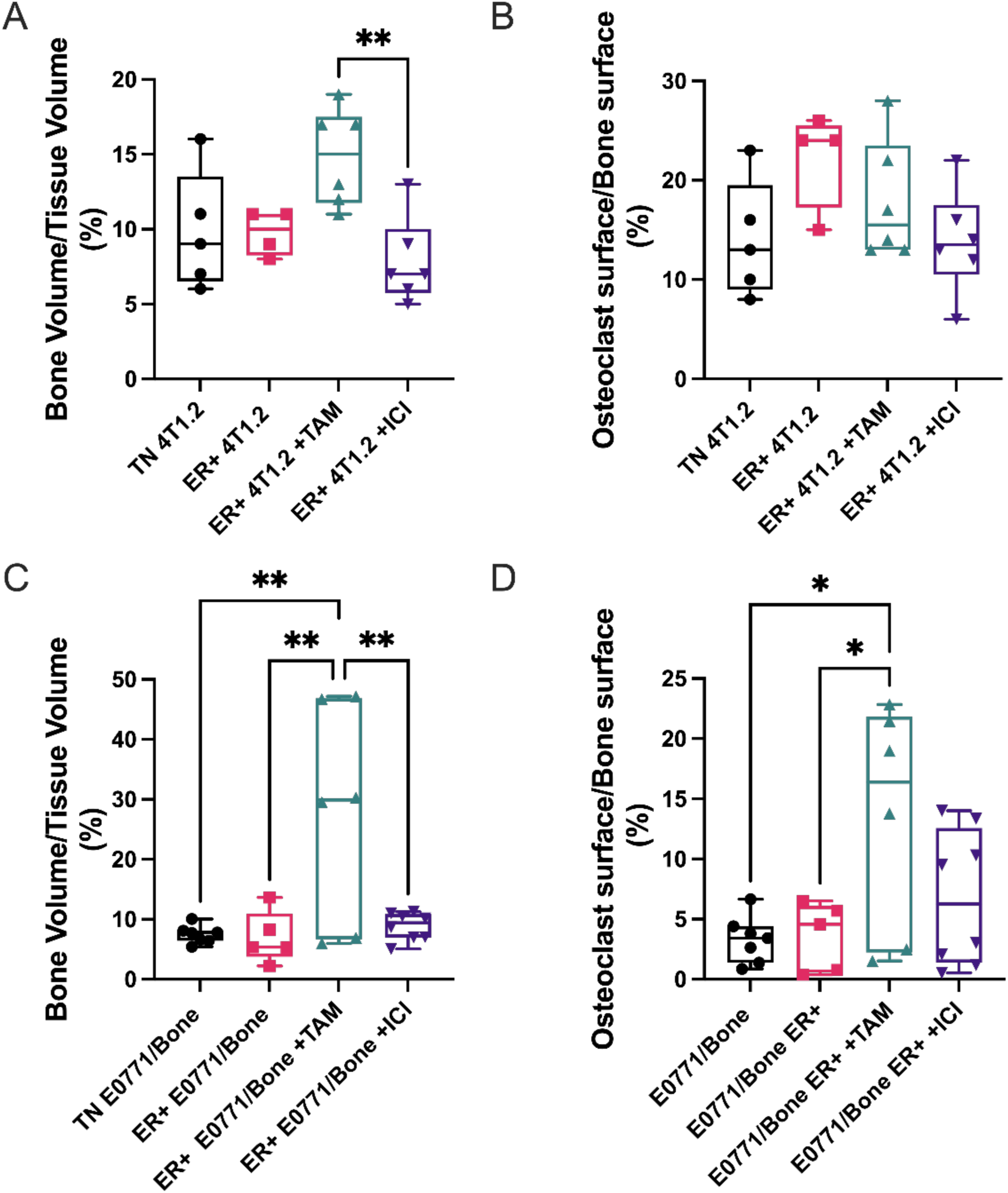
Tamoxifen induced alterations in the bone niche. Tibiae were isolated from mice after tumor implantation with 4T1.2 (A and B) or E0771/Bone (C and D) TNBC or ER+ breast cancer cells and treatment with vehicle controls, 5 mg/ 30-day time-release pellet tamoxifen (TAM), or 1 mg/wk ICI 182,780 (ICI). Bones were stained for TRAP+ osteoclasts and bone histomorphometry performed to calculate bone volume: tissue volume fraction (A and C) or Osteoclast Surface per Bone Surface (B and D). A significant decrease in bone volume: tissue volume was appreciated in the mice with 4T1.2 ER+ treated with ICI when compared with TAM (A). In both cell types, TAM treatment tended to have increased bone volume: tissue volume compared with all other groups (A and C). Mice with E0771/Bone ER+ tumors treated with TAM had significantly increased percentages of osteoclasts at the bone surface compared with ER+ E0717/bone without treatment and the E0771/Bone tumors (D). Both are represented as mean percentage ± SEM (n=4-8). * represents p<0.05 and ** represents p<0.01 by one-way ANOVA.

BC bone metastases are often osteolytic and activation of osteoclasts is associated with reactivation of dormant BC cells in the bone [31]. TRAP+ osteoclasts were quantified in mice implanted with ER+ and TN BC. No significant changes in osteoclast number were seen between ER+ and TN BC-injected mice (Figure 8B and D). ER+ E0771/Bone tumors treated with TAM had significant increased percentage of osteoclasts at the bone surface over total bone surface when compared with both ER+ E0771/Bone non-treated mice and with the parental E0771/Bone mice (Two-way ANOVA, p<0.05, Figure 8D). There were no significant differences in percentage of osteoclast surface to bone surface in any of the mice with 4T1.2 tumors (Figure 8B). Thus, neither osteoclast activation nor alterations in bone structure were responsible for the increased ER+ 4T1.2 bone metastasis.

## 4. Discussion

Bone metastasis from ER+ BC is a significant clinical problem with few advancements in treatment or prevention in recent decades. Two TN, bone-tropic cell lines, 4T1.2 and E0771/Bone which are derived from mouse mammary tumors and devoid of hormone receptors or human epidermal growth factor receptor 2, were transduced to express ERα. Through this we were able to produce ER+ cell lines that, when injected into the mammary glands of mice, established tumors that were responsive to two current antiestrogen therapies. The ER+ 4T1.2 cell line, unlike its TN parental line, metastasized to bone. ER+ tumors in the mammary glands not responded to antiestrogen treatment, but they had different immune profiles within the mammary gland with macrophages, B cells, and neutrophils being significantly increased or tending to be increased in the ER+ compared to TN tumors. The 4T1.2 ER+ model, which exemplifies important features of ER+ BC bone metastasis in women, will allow for further investigation into the initial steps, and potential treatment, of metastasis and preferential metastasis to bone of ER+ BC.

In our study, ER+ tumors were larger than TN tumors for both cell lines, indicating that ER within the cells was acting in a pro-tumorigenic fashion. When ER+ were treated with antiestrogen compounds, ER+ tumor weights and volumes decreased to approximately the size of the TN tumor, demonstrating that the transduced ER was functional and responsive to therapies targeted towards it. While this demonstrates that the tumor expresses a functional ER that these antiestrogen compounds can target, there were no differences in estrogen-related gene sets between the TN and ER+ tumors derived from the 4T1.2 cell line. Although unclear, this may be due to the fact that RNA sequencing of the tumors included not only tumor cells, but stroma, fibroblasts, inflammatory cells, and many other cell types that express ER [32]. It is possible that the addition of ER within the ER+ tumors did not overshadow the large amount of ER already present within the tumor microenvironment.

As mentioned above, RNA sequencing data from the TN and ER+ 4T1.2 mammary tumors did not show differences in estrogen receptor associated pathways. There were, however, significant downregulation of the RNA expression of genes associated with T cell receptor pathways in the ER+ 4T1.2 derived tumors. Also of note, CD3+ T cells tended to be decreased in ER+ tumors compared with TN tumors in our study. Decreased tumor infiltrating lymphocyte numbers has been observed in patients with ER+ disease when compared with TN tumors [33–36]. Additionally, of note is the set of 15 genes significantly downregulated in ER+ tumors in both humans and mice compared with ER- and TN tumors, respectively. Of those 15 shared genes, expression levels of 4 were associated with significant changes in patient survival, CARD11, ZAP70, GRAP2, and CD3G. Decreased expression of ZAP70, GRAP2, and CD3G is associated with worse survival in humans. This finding increases translatability and accuracy of disease modeling in mice and may provide for a novel model to guide novel, targeted therapies.

Aside from the influence of T cell infiltration into the mammary tumor, the influence of other immune cells within the tumor microenvironment has substantial impact on the prognosis of women. For example, tumor associated macrophages are associated with worse prognosis [27]. In our study, the mice with ER+ E0771/Bone derived tumors had significantly higher CD68+ macrophages and neutrophils within the tumor compared with the TN E0771/Bone derived tumors. Neutrophil infiltration is associated with increased metastatic spread, as neutrophil depletion prevents metastatic outgrowth [37] and colonization of metastatic niches [38]. Treatment of ER+ BC with SERMs, such as TAM, reduced T cell cytotoxicity while increasing neutrophil inflammation. SERDs, including ICI, increased infiltration of T cells [39]. Thus, antiestrogen treatment may further impact the immune landscape in ways that can be better studied with immunocompetent models. While assessing long-term disease progression was outside of the scope of our study, it is plausible that the differing immune landscapes seen in ER+ tumors when compared with TN may have implications for tumor progression.

A main purpose of our study was to develop an immunocompetent ER+ bone metastasis model. While we accomplished this, only mice with tumors from ER+ 4T1.2 cells, but not the ER+ E0771/Bone cells, developed metastases to the bone. Tumors in the ER+ 4T1.2 injected mice, but not TN, were present by 4 to 5 weeks post injection of transduced cells into the mammary glands. It is possible that with resection of the primary tumor and additional time, additional bone metastases would develop in more animals and possibly in the ER+ E0771/Bone model. Our short, 4-5 week time course for bone metastasis development is similar to the time course described in the seminal intra-mammary model of 4T1.2 cells, where cancer cells with stem-like properties were identified as early as 22 days following intra-mammary injection [40]. One major difference between our study and the previous report is that they used TN 4T1.2 cell lines to establish their models. Within a similar timeline, the previous study observed metastases from the TN 4T1.2-derived tumors, where we did not. This discrepancy may be due to differences in detection methods, with the previous study using a bone crush and clonogenic growth method and ours using histology, or which bones were isolated, the previous study focusing largely on the vertebral column and us focusing on the tibiae. Another model of intra-mammary injection of tumor cells with bone metastases found that bone metastasis did not occur within the first 30 days following injection of TN 4T1 tumor cells [41]. The xenograft model of ER+ ZR751 intraductal injection resulted in bone metastasis around day 88 [17], which is a much longer time course. While this may be due to differences in detection methods, as these studies used a luciferase reporting system to identify metastases, which may not be able to detect metastases if the cell numbers are small, it may represent differences in bone tropism and upregulation of metastatic mechanisms in the ER+ 4T1.2 cell line.

Interestingly, in our study all groups with tumors from ER+ 4T1.2 cells, regardless of antiestrogen therapy, had some evidence of bone metastasis. It is possible that these metastases occurred before the antiestrogen therapy began, since tumors were allowed to establish to 100 mm3 before treatment began. One of the antiestrogen compounds used in our study, TAM, is a SERM and is one of the most frequently used treatments for ER+ BC [42]. As a SERM, TAM works in an antiestrogen manner in the tumor to inhibit tumor growth, while acting in a pro-estrogen manner in the bone to maintain bone volume in post-menopausal women [42,43]. The result reported here are consistent with previous studies; mice treated with TAM had increased bone volume percentages when compared with mice treated with a vehicle control. It is important to note that mice with TN tumors were not treated with antiestrogen compounds in this study, so the influence of TAM on the bone microenvironment, although likely similar to the response in the mice with ER+ tumors, could not be assessed.

We did identify other changes in the bone microenvironment between mice with TN and ER+ tumors without evidence of bone metastasis, thus representing a premetastatic niche. The percentages of Sca-1 positive and endomucin vasculature, markers for hematopoietic stem/tumorigenic cells and blood vessels, respectively, within the bones of mice with ER+ E0771/Bone derived tumors were significantly increased when compared with TN derived tumors. Previous studies across multiple cancer types have demonstrated that tumors can modulate the premetastatic bone microenvironment to become a more hospitable environment by increasing hematopoietic stem cells and vasculature [44–46]. Thus, the numbers of potential colonization niches were increased in mice with ER+ E0771/Bone tumors. While the specific mechanisms driving the premetastatic increase in stem cell and endothelial cells is unclear in our study, it is apparent that the expression of ER within the E0771/Bone, but not the 4T1.2, mammary tumors were associated with downstream modification of the bone microenvironment.

While the 4T1.2-based model may be useful in the future to study ER+ BC bone metastasis, several shortcomings of the study must be acknowledged. Relatively few mice were used in each cohort and relatively few mice with ER+ BC developed bone metastasis. The study was designed and powered to measure changes in the primary tumor as the rate of metastasis was initially unknown. Since monitoring for bone metastasis was done by histological and immunohistochemical evaluation of the tibia, it is possible that some bone metastases were missed. Future studies will include the introduction of luciferase into the cells. Initial characterization was performed without luciferase as its introduction into 4T1 cells was shown to reduce metastatic spread due to interaction with the immune system [47]. Furthermore, RNA sequencing on the metastasis was not performed which could give important information into the characteristics of the metastatic population of cells versus the primary tumor. Lastly, mice with TN tumors were not treated with antiestrogen compounds. While there we would not expect tumor volumes and weights in the TN group to change since there is not a functioning ER, these treatments will likely alter the bone microenvironment and could affect metastasis and immune cell infiltration.

## 5. Conclusions

In summary and conclusion, we generated an immunocompetent model of ER+ BC that spontaneously metastasizes to bone with the ER+ 4T1.2 cells. This model allows for further exploration of bone metastasis mechanisms and for the development of new therapeutics in the presence of an intact immune system, which may translate into improved clinical outcomes for women with bone metastasis from ER+ BC.

## AUTHOR CONTRIBUTIONS

Conceptualization, RS, KLC, and BAK; methodology, KLL, LS, ASW, KLC, and BAK; formal analysis, KLL, BW, LS; investigation, MTX, JT, VES; writing—original draft preparation, KLL and LS; writing—review and editing, KLL, LS, ASW, BW, MTX, VES, JT, RS, KLC, and BAK; visualization, KLL, LS, BW; supervision, RS, KLC and BAK; funding acquisition, KLC and BAK. All authors have read and agreed to the published version of the manuscript.

## DATA AVAILABILITY STATEMENT

RNA sequencing data will be made publicly available at GEO; a reference will be provided when in review.

## ACKNOWLEDGEMENTS

We would like to thank Dr. Elaine Alarid for the generous donation of the ERα-GFP construct. The authors would also like to thank Amanda Salis at the Salis Institute for her time and effort editing the manuscript.

